# The extended default mode network dysconnectivity in schizophrenia

**DOI:** 10.64898/2026.11.10.765658

**Authors:** Tahereh S. Zarghami

## Abstract

The default mode network (DMN) extends beyond cortex into subcortical structures, yet how this extended DMN (eDMN) is causally reorganized in schizophrenia (SZ) remains unclear. Here, resting-state fMRI from 74 healthy controls (HC) and 66 SZ patients was analyzed. Subject-specific eDMNs were identified using spatially-constrained independent component analysis, and effective connectivity among 13 regions was estimated with spectral dynamic causal modeling, with group effects assessed using parametric empirical Bayes. Causal centrality—a dynamics-sensitive and causality-based measure of nodal importance—was computed per region, compared between groups, and correlated with MATRICS cognitive scores in patients. Causal centrality rankings were highly similar across groups (ρ = 0.87), with both favoring subcortical over cortical regions. Group differences were regionally specific: temporal cortex was more causally central in patients, whereas posterior parietal cortex was more causally central in controls. Most connection-level SZ−HC differences originated from ventrolateral prefrontal cortex, prefrontal cortex, basal forebrain, temporal cortex, and thalamus. In patients, temporal causal centrality correlated negatively with reasoning/problem-solving, and ventrolateral prefrontal causal centrality with processing speed and composite cognition. The eDMN’s core causal architecture is preserved in schizophrenia, but temporal influence is enhanced and parietal influence attenuated— alterations that are behaviorally meaningful and consistent with pathological reorganization.

## Introduction

The default mode network (Buckner & DiNicola, 2019; Raichle et al., 2001) is a collection of brain region that are typically deactivated during external goal-directed tasks, but activated during internally focused cognitive processes such as mind wandering, memory retrieval, self-referential and emotional processing, perspective taking, etc. (Knyazev, Savostyanov, Bocharov, Levin, & Rudych, 2020; Murphy et al., 2018; Mwilambwe-Tshilobo & Spreng, 2021; Qin & Northoff, 2011). Most of these psychological functions are readily carried out during wakeful *resting state*, the context in which DMN was first characterized^1^ (Greicius, Krasnow, Reiss, & Menon, 2003; Raichle et al., 2001).

To date, DMN has largely been a cortically defined network, consisting of regions distributed across the ventromedial and lateral prefrontal cortex, posteromedial and inferior parietal cortex, as well as the lateral and medial temporal cortex. While DMN has been considered a backbone of cortical integration (Alves et al., 2019; Andrews-Hanna, Reidler, Sepulcre, Poulin, & Buckner, 2010; Kernbach et al., 2018; Lopez-Persem, Verhagen, Amiez, Petrides, & Sallet, 2019; Margulies et al., 2016), the involvement of subcortical regions in the DMN has been usually overlooked. Importantly, subcortical structures contain the neurochemically diverse nuclei that are crucial in the pathophysiology of various brain diseases in which DMN connectivity is gravely affected; these include Alzheimer’s disease, Parkinson’s disease, schizophrenia, depression and temporal lobe epilepsy, among others (Kottaram et al., 2019; Mothersill et al., 2017; Parsons, Bowden, Vogrin, & D’Souza, 2020; Qian et al., 2019; Ruppert et al., 2021; Sheline et al., 2009; Whitfield-Gabrieli & Ford, 2012). Therefore, investigating the contribution of subcortical structures to the DMN function is an essential step towards understanding the mechanisms of DMN disruption in these disorders, and more broadly serves to reconcile neurochemistry, connectivity and cognition.

More recently, (Alves et al., 2019) have revisited the spatial extent of DMN, with specific focus on the structural and functional role of the subcortical regions in the DMN integration. The result has been an improved neuroanatomical model of the DMN, consisting of 33 cortical and subcortical regions of interest (ROIs). These regions and their MNI coordinates have been outlined in Table 2 of (Alves et al., 2019), and the final template has been shared on the NeuroVault^2^ repository.

In previous work (Zarghami, 2023), we used this extended DMN (eDMN) as prior spatial information to identify subject-specific eDMNs in 74 healthy subjects, which were then causally modeled using spectral DCM (Friston, Kahan, Biswal, & Razi, 2014; Razi, Kahan, Rees, & Friston, 2015). Thereafter, the group-average causal structure of eDMN was characterized using parametric empirical Bayes. We then argued that classic graph theoretical centralities are not apt for quantifying the importance of nodes/regions in dynamic causal models, since they are “poor substitutes” for *causal influence* of the nodes (Dablander & Hinne, 2019) and do not account for the specific “flow process”, i.e. *dynamics*, on the graph (Borgatti, 2005). And, the *cyclic* nature of DCM poses further challenges for quantifying the influence of the nodes. To address these issues, we introduced *causal centrality*, which can gauge the importance of nodes in a dynamic causal model (i.e. a directed causal graph) using the notion of *intervention* on graphical models.

The *causal centrality* of a node in a DCM was defined (Zarghami, 2023) as the normalized KL- divergence between the pre- and post-intervention distributions of the DCM; where the distribution is the joint probability distribution over the model’s unobservable^3^ (*θ*) and observable (*Y*) variables, given the generative model of DCM (*m*); and intervention refers to removing the causal influences of that node directed at the other (hidden) nodes.

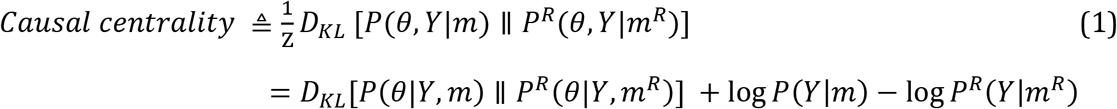

where *P* and *P^R^* refer to the pre- and post-intervention distributions, associated with the *full* and *reduced* models (*m* and *m^R^*); and the normalization constant *Z* is the model evidence. In the reduced model (*m^R^*), some of the edges (*θ_i_*) of the full model have been removed; i.e., forcefully set to zero. As previously shown (Zarghami, 2023), this expression is *identifiable*, using *Bayesian model reduction* (BMR) (Friston et al., 2016; Friston, Parr, & Zeidman, 2018; Friston & Penny, 2011), which facilitates the computation of post-intervention distributions from pre-intervention quantities. Specifically, when the inference is variational, the causal centrality expression simplifies to:

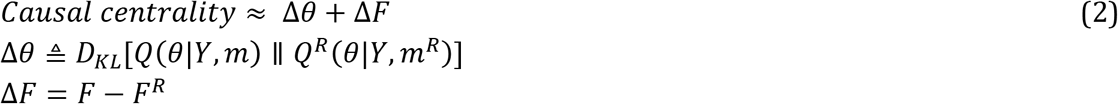

The approximate posterior (*Q*(*θ*|*Y*, *m*) ≈ *P*(*θ*|*Y*, *m*)) and the free energy^4^ (*F* ≈ log *P*(*Y*|*m*)) of the full model are already available by virtue of the original model inversion. Hence, BMR can analytically compute *Q^R^* and *F^R^* for the reduced (intervened upon) model. Eq. 2 further reveals the intuitive properties of a node with high causal centrality: intervention on the causal effects of a highly central node would result in substantial drop^5^ in model evidence (Δ*F* = *F* − *F^R^* ≫ 0), and the remaining causal strengths should be substantially revised to explain the same data *Y* (Δ*θ* ≫ 0).

We already provided face, construct and empirical validity for this measure for a group of healthy subjects in previous work (Zarghami, 2023). In the present study, we extend the analysis to a cohort of schizophrenic patients. Specifically, we intend to investigate the extent to which our original findings—about the critical role of subcortical structures in the causal backbone of eDMN in HC subjects—holds in the SZ group. We also intend to quantify potential differences in the effective connectivity and causal centralities of eDMN regions between the two groups. Whether nodal importance—in terms of causal centrality—is related to behavioral (cognitive) measures in the patient group is another question that we query this dataset for.

## Materials and Methods

In this section, the causal architecture of the extended DMN is identified in a group of healthy and schizophrenic subjects; the commonalities and differences in effective connectivity of the two groups are quantified. Then, causal centralities of the constituent regions are computed for each group and contrasted against each other, while the overall centrality *ranking* of the regions are compared across groups. Finally, linear relation between the causal centrality of selected regions and the cognitive scores of the subjects is investigated in the patient group.

### Dataset and pre-processing

We analyzed the publicly available schizophrenia dataset of the Center for Biomedical Research Excellence (COBRE)^6^ (Çetin et al., 2014), which includes 72 schizophrenic (SZ) and 76 healthy control (HC) subjects (18-65 years old). The patients had been diagnosed using the Structured Clinical Interview for DSM-IV^7^ Axis I Disorders (SCID-I) (First, Spitzer, Gibon, & Williams, 2002), and were receiving antipsychotic medications. Five-minute resting state scans were acquired on a 3-Tesla Siemens Tim Trio scanner, during which subjects were instructed to keep their eyes open and fixate on a central cross. A total of 150 (T2*-weighted) functional volumes were collected using a gradient-echo EPI sequence, with the following settings: TR = 2 s, TE = 29 ms, flip angle = 75°, 33 axial slices, ascending acquisition, matrix size = 64 × 64, voxel size = 3.75 × 3.75 × 4.55 mm, field of view = 240 mm. A high-resolution T1-weighted structural image had also been collected for each subject.

Standard pre-processing of functional data (before independent component analysis, ICA) was performed using the SPM12 software^8^. In brief, the first 5 volumes were discarded to allow for T1 equilibration; the remaining images were realigned to the first volume (for motion correction), slice-timing corrected, co-registered to the anatomical image of the corresponding subject, warped to the standard Montreal Neurological Institute (MNI) template (Collins et al., 1998), resampled to 3 *mm*^3^ isotropic voxels, and smoothed with a Gaussian kernel (FWHM = 6 mm).

Two disenrolled subjects, one subject with incomplete functional session and five subjects with excessive head motion (translation > 4 mm or rotation > 4 degrees) were excluded. For the remaining subjects, the mean framewise displacement (FD) was reasonably low (0.41 ± 0.22 mm) (Power, Barnes, Snyder, Schlaggar, & Petersen, 2012). The demographics of the retained 140 subjects (74 HC/66 SZ) have been summarized in Table 1. There were no significant differences in age, gender proportion, maximum head translation and rotation between the two groups (uncorrected p-values = 0.12, 0.26, 0.13, 0.36, respectively). The patients were receiving antipsychotic medications, which were converted to their chlorpromazine (CPZ) equivalents.

**Table 1:**
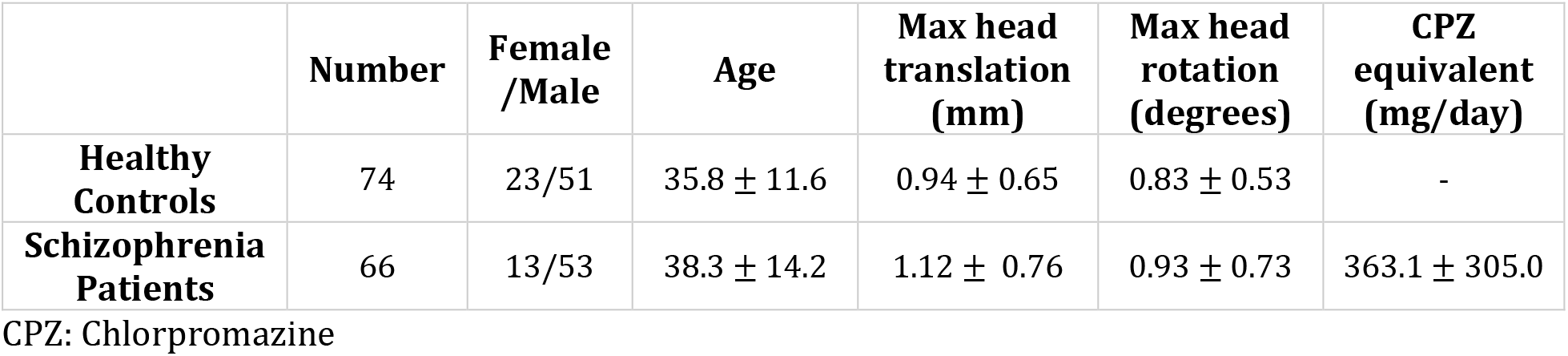
Demographics of the participants.

**Table 2:** The regions of eDMN included in DCM analysis.

| **Abbreviation** | **Region** |
|---|---|
| **PFC** | Prefrontal cortex |
| **Tmp** | Temporal cortex |
| **PCC** | Posterior cingulate cortex |
| **Rsp** | Retrosplenial cortex |
| **PPC** | Posterior parietal cortex |
| **VLPFC** | Ventrolateral prefrontal cortex |
| **Amy** | Amygdala |
| **BF** | Basal forebrain |
| **C** | Caudate |
| **Cb** | Cerebellar region |
| **PH** | Parahippocampal region |
| **T** | Thalamus |
| **MidB** | Midbrain |

The MATRICS^9^ consensus cognitive battery (MCCB) tests (August, Kiwanuka, McMahon, & Gold, 2012; Kern et al., 2011; Nuechterlein et al., 2008) had been conducted to estimate the cognitive performance of the participants in seven domains: speed of processing, attention/vigilance, working memory, verbal learning, visual learning, reasoning/problem solving, and social cognition. The composite MCCB score had been calculated as the standardized sum of all seven domains, based upon published normative data (Kern et al., 2011).

### Identifying eDMN

To identify the extended DMN for each subject, the eDMN template (Alves et al., 2019) was used as prior information in a spatially-constrained independent component analysis (SC- ICA) (Lin, Liu, Zheng, Liang, & Calhoun, 2010). Constrained ICA is a semi-blind source separation method, where prior information is added to the contrast function of a standard blind ICA in the form of (in)equality constraints. As such, the prior information ensures some level of similarity between the recovered sources and the reference templates, and keeps the optimization solution bounded. Importantly, SC-ICA accommodates the spatial variability of the networks between subjects (unlike fixed templates) while maintaining the component (i.e., network) correspondence over the group. SC-ICA algorithm has been implemented in the Group ICA of fMRI Toolbox (GIFT^10^). The template from (Alves et al., 2019) and the average spatial map of eDMN from the present study have been illustrated in Fig. S1 and Fig. S2, respectively.

### Region specification and time series extraction

Several eDMN regions were merged based on physical and functional proximity for computational tractability of the DCM analysis^11^. First, the 16 pairs of highly correlated homologous regions were merged. Next, VMPFC^12^, AMPFC^13^ and DPFC^14^ were grouped into one prefrontal (PFC) component; two temporal sub-regions (TP^15^ and MTG^16^) were merged into one temporal (Tmp) component; and two cerebellar regions (CbH^17^ and CbT^18^) were represented by one cerebellar (Cb) component. This resulted in 13 regions for spectral dynamic causal modeling, which have been listed in Table 2.

The average regional time series were extracted for these 13 ROIs from the intersection of group-adjusted individual eDMNs with the parcellation map of (Alves et al., 2019). Post- processing (Allen et al., 2014; Damaraju et al., 2014; Iraji et al., 2022) of the time series included: removal of linear trends, regression of nuisance (motion and physiological) signals, and replacement of outlier time points with spline interpolations (using AFNI’s 3dDespike). The average functional connectivity between these regions has been depicted in Fig. S3. The post-processed time series were next modeled using spectral DCM.

### Spectral dynamic causal model

Spectral DCM was used to estimate the effective (causal) connections between the 13 regions of eDMN, for each subject. The generative model of this DCM specifies how the observed complex cross-spectra^19^ of fMRI signals are generated from the interaction of coupled neuronal ensembles and the neurovascular coupling. The model is initially expressed in the time domain as a state-space model with two equations; the first equation describes the dynamics of an endogenously driven network of coupled neuronal populations, and the second equation maps these neuronal activities to the observed hemodynamic responses (Friston et al., 2014; Razi et al., 2015):

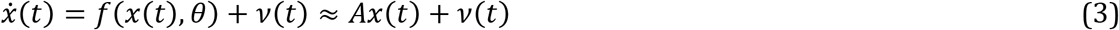

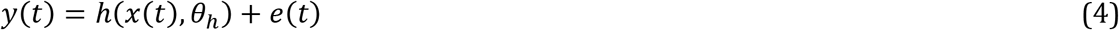

where *x*(*t*) is a vector containing the hidden neural states (i.e., ensemble activities) of *n* brain regions, at time *t*. The rates of change^20^ in neural states *x͘*(*t*) are influenced by the activities of (self and) other regions via the connectivity matrix *A*, and fast endogenous neural fluctuations *ν*(*t*). The hemodynamic response function ℎ (Stephan, Weiskopf, Drysdale, Robinson, & Friston, 2007) translates the neural activities into measured BOLD^21^ signals *y*(*t*), which also contain the observation noise *e*(*t*). After linearization, Fourier transformation and parametrization of the noise cross-spectra, this generative model admits a deterministic form that can be efficiently *inverted* using variational Bayesian methods (Friston et al., 2007; Friston et al., 2014; Razi et al., 2015).

Model inversion refers to the estimation of optimal model parameters, given a generative model and some observations. In variational Bayesian inference, this is achieved by maximizing free energy (*F*, a lower bound on log model evidence) with respect to the approximate posteriors of the parameters, *Q*(*θ*|*y*, *m*) (Friston et al., 2007; Zeidman et al., 2022). Notably, free energy offers a trade-off between model accuracy and complexity (F = accuracy – complexity), with the latter term protecting against overfitting. Moreover, maximized free energy serves as a proxy for log model evidence (*F_opt_*(*Q*) ≈ log *P*(*y*|*m*)), which is used to compare the plausibility of different models explaining the same data. The associated optimized posteriors (*Q_opt_*(*θ*|*y*, *m*) ≈ *P*(*θ*|*y*, *m*)) are used to perform inference on the parameters. The most prominent parameters of DCM are the effective connections encoded in the *A_n_*_×*n*_ matrix. Each off-diagonal entry *a_ij_* of this matrix denotes the causal influence of region *j* on region *i*, which can be excitatory (positive) or inhibitory (negative). Conversely, the self-connections^22^ (*a_ii_*) are parametrized with negativity constraints to ensure dynamical stability of the model (Zeidman, Jafarian, Corbin et al., 2019).

This concludes our brief description of the generative model of spectral DCM and the associated Bayesian model inversion scheme. Having inverted each subject’s DCM, group analysis was conducted using a hierarchical Bayesian framework known as parametric empirical Bayes (PEB)—as elaborated next.

### Group analysis using parametric empirical Bayes

PEB is a hierarchical Bayesian model, particularly useful for estimating group effects in DCM studies. Operationally, PEB is a Bayesian general linear model (GLM) that partitions between-subject variability into certain designed group effects (such as group mean and difference) and some additive random effects. In contrast to the *summary statistic* random effects approach based on point estimates, PEB accommodates the *full posterior* densities of the parameters. In its simplest form, PEB can be expressed as:

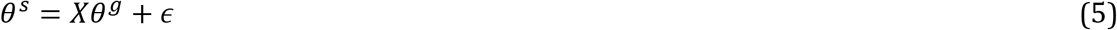

where *θ^s^* includes the posteriors of *p* DCM parameters for *N* subjects; *X_Np_*_×*pd*_ is the design matrix for *d* regressors replicated over *p* parameters; *θ^g^* denotes the group parameters; and *ε_Np_*_×1_ contains the random effects. The columns of X encode the hypothesized sources of inter-subject variability (e.g., group mean). To invert a PEB model, subject- and group-level parameters are estimated iteratively in a variational Bayesian scheme (Friston et al., 2007; Friston et al., 2016; Friston, Zeidman, & Litvak, 2015; Zeidman, Jafarian, Seghier et al., 2019).

In the present study, PEB was used to compute the group mean value for each effective connectivity parameter, per group. In the SZ group, in addition to age and gender, the group effects were adjusted for medication dosage as well. These PEB (i.e., group DCM) posteriors comprised the directed cyclic graph for which causal centralities of the nodes were computed. Two additional PEB analyses were performed to compute the global mean (i.e. commonality) of the two groups and their difference (SZ-HC contrast). Each PEB structure was optimized post-hoc using exploratory BMR (Friston et al., 2016; Friston & Penny, 2011). We will now go over the main results.

## Results

### Causal structure of the eDMN

Spectral DCM was used to estimate the effective (causal) connections between the regions of eDMN (Table 2), for each subject. DCM model fitting was assessed using the coefficient of determination, *R*^2^, which reflects the proportion of variance in the observations (i.e., cross- spectra) that is explained by the spectral DCM model^23^ (*R*^2^ = 0.91 ± 0.03). Group analysis was conducted in the PEB framework.

Group-specific connectivities^24^ have been plotted in Fig. 1, while the global average and group difference patterns are demonstrated in Fig. 2. Overall, these plots suggest that the causal influences among the cortical regions of eDMN are dense and mostly excitatory. Conversely, within-subcortical causal architecture seems more balanced in terms of excitatory and inhibitory effects. Notably, the cortical-to-subcortical causal influences are mostly excitatory, whereas the subcortical-to-cortical effects are more inhibitory. In Figs. S4-S7 the posterior *uncertainties* of the group effective connections have been illustrated in addition to the posterior expectations.

**Fig. 1:**
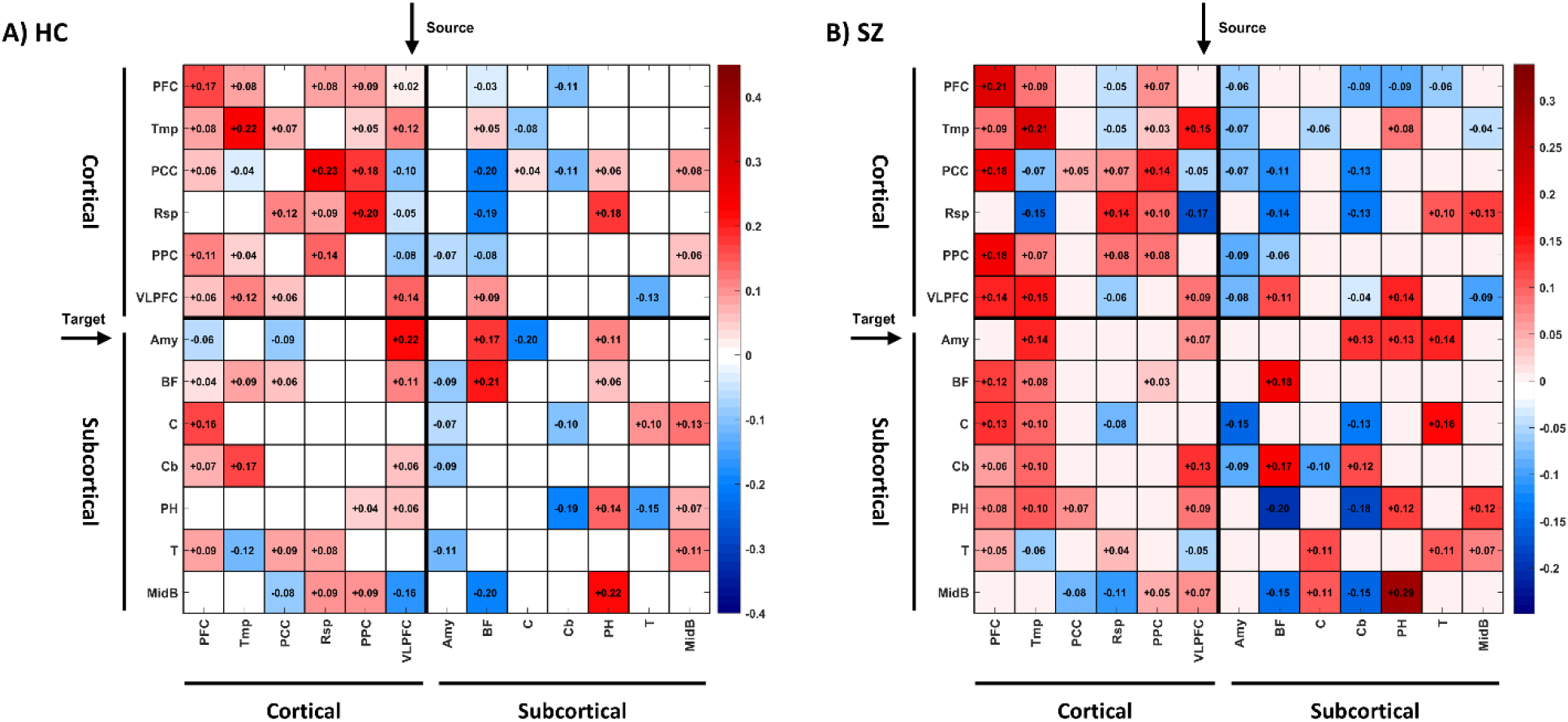
Group-average DCM results. A) HC group B) SZ group. Columns and rows denote source and target regions, respectively. The expected connection strengths have been color-coded, where red and blue indicate excitatory and inhibitory effects, respectively. The regions have been grouped to cortical and subcortical, for clarity. The posterior uncertainties of the connections have been depicted in supplementary Fig. S4 and Fig. S5.

**Fig. 2:**
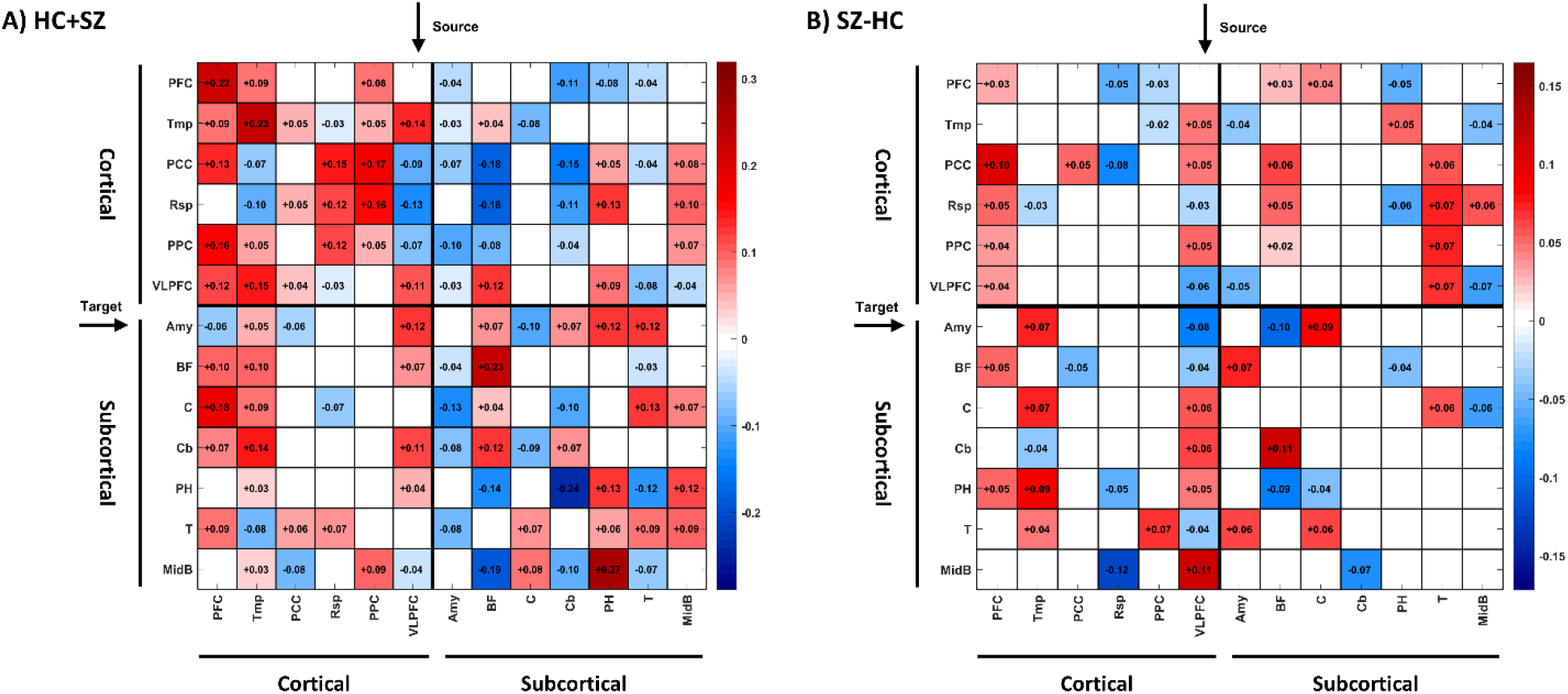
Group mean and difference DCM results. A) Global mean effect across HC and SZ subjects B) SZ-HC effect. Columns and rows denote source and target regions, respectively. Red and blue indicate excitatory and inhibitory effects, respectively. The posterior uncertainties of the connections have been depicted in supplementary Fig. S6 and Fig. S7.

Notably, in the group-difference graph (Fig. 2-B), the causal influences from VLPFC, PFC, BF, Tmp and thalamus to the rest of the regions bear most of the difference effects. We will go over to the implications of these findings in the Discussion. After estimating the average causal influences among the eDMN regions, the node centralities were computed based on group-specific causal graphs (in Fig. 1), as elaborated next.

### Centrality of the eDMN regions

To identify the nodes whose overall causal influence differs in the eDMN of HC and SZ subjects, causal centrality (Eq. 2) was computed for each node and subject, and the values were compared between the two groups using permutation-based two-sample t-tests, followed by false discovery rate (FDR) correction. Moreover, Spearman’s rank correlation was used to quantify the similarity in the average causal centrality ranking of the nodes, in the HC and SZ groups, which are listed in Table 3. Rank correlation turned out to be 0.87 (p- value = 1.25e-4), suggesting that the average causal centrality ranking is highly similar in the two groups. In supplementary Fig. S9 we have shown that the causal centrality ranking in the SZ group shows a trend of anti-correlation with graph theoretical centrality rankings, similar to the results obtained previously in the HC group (Fig. S8) (Zarghami, 2023). That is, subcortical regions get higher ranks in terms of causal centrality but lower ranks based on certain graph theoretical centralities. The reverse is true for cortical regions.

**Table 3:** Causal centrality scores and ranks of the eDMN regions, based on group-DCM (i.e. PEB) results. Higher ranks correspond to higher centrality values. The maximum/best rank is 13, which is the number of modeled regions, as listed in Table 2.

|  | **Causal Centrality** |  |  |  |
|---|---|---|---|---|
|  | **HC** |  | **SZ** |  |
| **Region** | **Value** | **Rank** | **Value** | **Rank** |
| **PFC** | 2626.35 | 1 | 2636.78 | 1 |
| **Tmp** | 3581.82 | 2 | 5022.93 | 2 |
| **PCC** | 6321.73 | 3 | 6546.78 | 4 |
| **Rsp** | 9708.33 | 7 | 10648.35 | 9 |
| **PPC** | 9305.69 | 6 | 7066.95 | 5 |
| **VLPFC** | 7878.70 | 5 | 7584.73 | 6 |
| **Amy** | 12085.05 | 11 | 13479.00 | 12 |
| **BF** | 6361.08 | 4 | 6161.73 | 3 |
| **C** | 10745.96 | 8 | 10571.41 | 8 |
| **Cb** | 12282.63 | 12 | 11170.84 | 10 |
| **PH** | 11663.28 | 10 | 11488.36 | 11 |
| **T** | 10930.42 | 9 | 8754.27 | 7 |
| **MidB** | 16030.32 | 13 | 14825.87 | 13 |

As for per-region comparison, permutation-based t-tests revealed that the average causal centrality of the Tmp region is significantly higher in the patient group (qFDR = 0.013, effect size = 0.57), whereas that of PPC is significantly higher in the HC group (qFDR=0.019, effect size = 0.49). The results have been illustrated in Fig. 3.

**Fig. 3:**
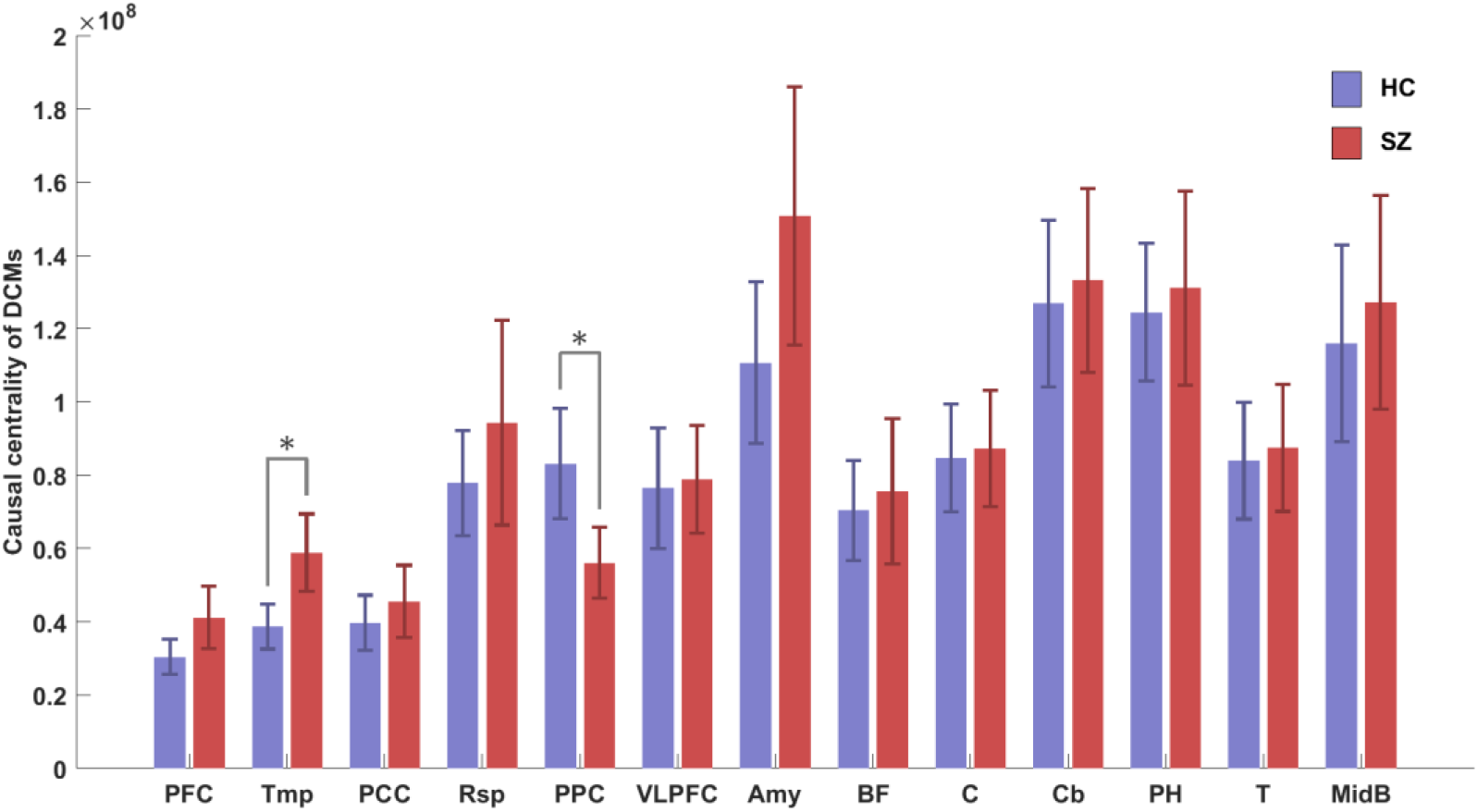
Comparison of average centrality of regions between HC and SZ groups. Tmp and PPC regions show significant differences. Bars indicate average causal centrality values; lines show 95% confidence intervals and * stands for qFDR < 0.05. Tmp: Temporal cortex. PPC: Posterior parietal cortex. Abbreviations are listed in Table 2.

### Cognitive relevance

The linear relationship between the causal centrality of influential regions and the MCCB cognitive scores of the SZ subjects was assessed using correlation analysis. The results showed that causal centrality of the Tmp region is negatively correlated with reasoning/problem solving capability of the patients (*ρ* = −0.37, *pvalue* = 0.003, *qFDR* = 0.025). Additionally, causal centrality of VLPFC is negatively correlated with the processing speed (*ρ* = −0.31, *pvalue* = 0.012, *qFDR* = 0.05) and composite MCCB scores (*ρ* =−0.34, *pvalue* = 0.008, *qFDR* = 0.05) of the patients. Fig. 4 includes the scatter plots as well as the linear fits.

**Fig. 4:**
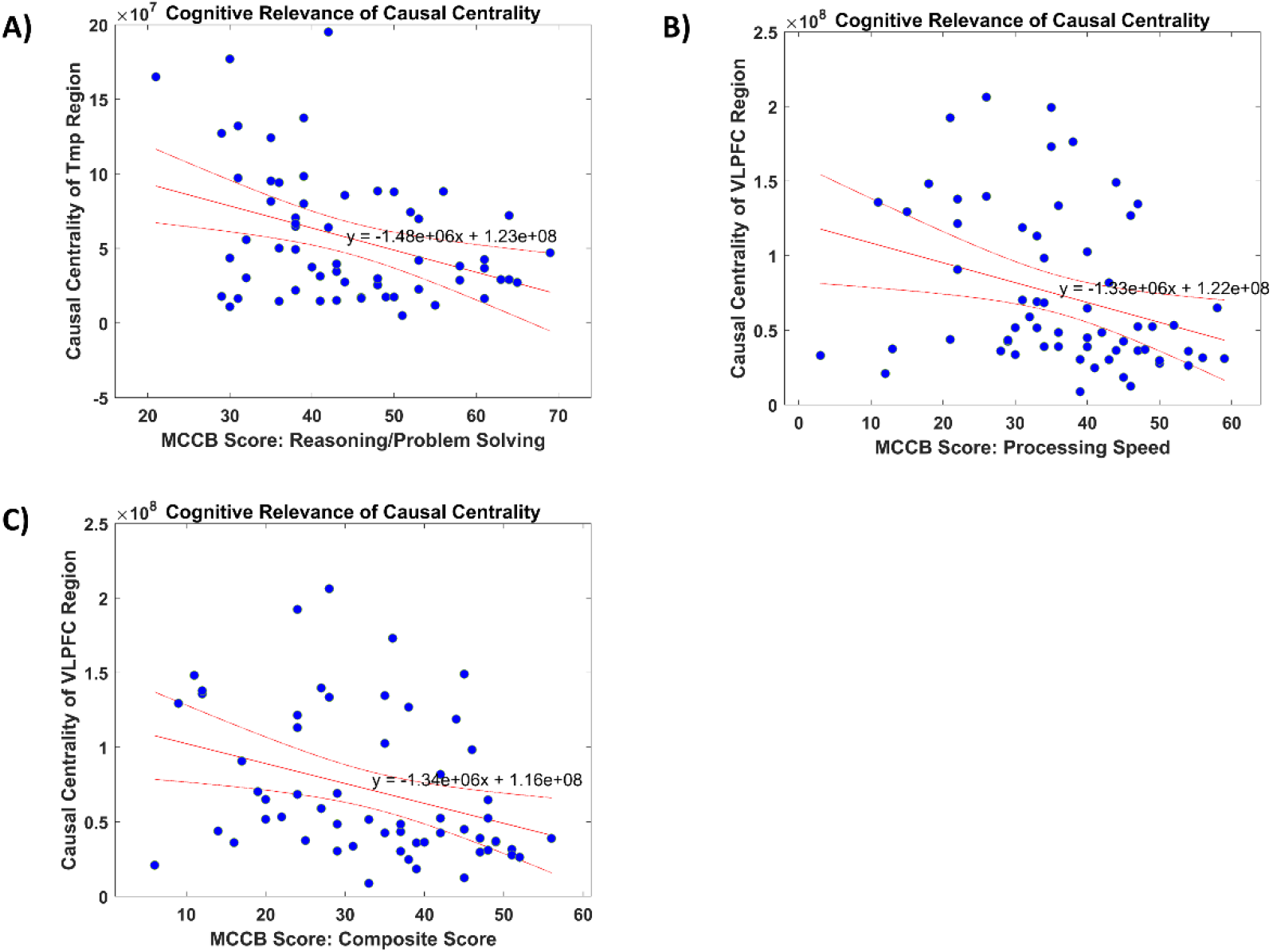
Behavioral (cognitive) relevance of regional causal centrality in the SZ group. A) Causal centrality of Tmp region is negatively correlated with reasoning/problem solving capability of the patients. B) Causal centrality of VLPFC is negatively correlated with processing speed of the patients. C) Causal centrality of VLPFC negatively correlates with composite MCCB scores of the patients. Correlation values are mentioned in the main text.

## Discussion

We characterized the extended DMN in HC and SZ subjects, in resting state, using spectral dynamic causal modeling and parametric empirical Bayes. Most of the effective connectivity differences originated from VLPFC, PFC, BF, Tmp and thalamus. Notably, causal centrality of VLPFC was negatively correlated with the processing speed and composite MCCB cognitive scores of the patients, while causal centrality of the Tmp region was negatively corelated with reasoning/problem solving capability of the patients. We will shortly elaborate on the significance of these regions in the pathophysiology of SZ.

Interestingly, the main composition of eDMN—in terms of causal centrality ranking of the regions—was highly similar in HC and SZ groups (rank correlation = 0.87), with both groups favoring subcortical regions over corticals. These results corroborate previous findings about the crucial role of subcortical structures in the neuromodulatory systems of the brain, and highlight their contribution to the organization of large-scale networks, in this case the eDMN (Horn et al., 2017; Horn et al., 2019; Horn & Fox, 2020; Li et al., 2021; Schiff et al., 2007; Zarghami, 2023)

Region by region comparison revealed that the Tmp region in the SZ group is more causally central compared to the Tmp region in the HC group. The reverse was true for the PPC region, which was more causally central in the HC group. Enhanced causal centrality of the Tmp cortex in the eDMN of SZ subjects is consistent with the presumed role of this region in dysfunctional auditory processing and auditory hallucinations of the patients (Hugdahl et al., 2008; Hugdahl, Løberg, & Nygå rd, 2009). On the other hand, suppressed causal centrality of PPC in the patient group can explain working memory deficits and cognitive impairment in SZ, given the implication of PPC in cognitive functions, especially working memory (Nielsen et al., 2017). To further put our findings in context, we review the role of the top regions— that drive group differences in our results—in the pathophysiology of SZ. These regions include: VLPFC, PFC, BF, Tmp, Thalamus and PPC.

The ventrolateral prefrontal cortex (VLPFC) supports working memory, response inhibition, and selection of task-relevant information, and its dysconnectivity is well documented in schizophrenia (Minzenberg, Laird, Thelen, Carter, & Glahn, 2009; Ragland et al., 2009; Ragland et al., 2015; Wolf et al., 2007). The present finding that VLPFC efferents contribute disproportionately to the group-difference graph is consistent with this literature. More striking is the negative correlation between VLPFC causal centrality and both processing speed and composite MCCB scores. These results could imply that in patients, the causal prominence of VLPFC is not compensatory but tracks with worse cognition. The network might have become increasingly reliant on a region whose dynamics are already compromised, yielding poorer behavioral output.

The broader prefrontal cortex (PFC), particularly medial and dorsomedial sectors, is central to the DMN’s internally directed cognition, and PFC dysconnectivity is among the most replicated findings in schizophrenia (Anticevic et al., 2015; Arnsten, 2011; Perlstein, Dixit, Carter, Noll, & Cohen, 2003; Zhou, Fan, Qiu, & Jiang, 2015). PFC connections were among the top contributors to between-group differences, consistent with the view that PFC is reorganized rather than merely hypofunctional in its causal architecture.

The basal forebrain (BF) cholinergic nuclei provide the principal cholinergic innervation to prefrontal cortex, hippocampus, and amygdala, and are critical for attention and memory (Ballinger, Ananth, Talmage, & Role, 2016; Mesulam, 2013), with altered BF cholinergic structure and receptor availability reported in psychotic disorders (Eickhoff et al., 2022; Saint-Georges et al., 2025; Scarr, Gibbons, Neo, Udawela, & Dean, 2013). That BF efferents contributed substantially to the SZ-HC difference graph is mechanistically plausible, given the BF’s position to modulate large-scale cortical dynamics. Importantly, BF causal centrality per se did not differ significantly—only its connections did. This distinction suggests preserved overall causal importance but altered projection targets or strengths, consistent with a rerouting rather than a loss of neuromodulatory control.

The temporal region (Tmp), particularly superior and middle temporal gyri, is implicated in auditory and language processing, and its dysfunction is a longstanding candidate mechanism for auditory hallucinations and thought disorder in schizophrenia (Hugdahl et al., 2008; Hugdahl, Løberg, & Nygå rd, 2009; Kaur et al., 2020; Lennox, Park, Medley, Morris, & Jones, 2000). Tmp causal centrality was significantly higher in patients and negatively correlated with reasoning/problem-solving scores. A temporal region that becomes more causally dominant within the eDMN may reflect increasing entrainment to temporally mediated processes at the expense of flexible reasoning; the negative correlation suggests this shift is not benign, possibly because the network is captured by temporal dynamics that interfere with prefrontal reasoning circuits.

The thalamus is a critical relay within cortico-striato-thalamo-cortical loops, with meta- analytic evidence for reduced thalamic volume and convergent findings of thalamocortical dysconnectivity in schizophrenia—hyperconnectivity with sensorimotor cortex and hypoconnectivity with prefrontal-limbic regions (Anticevic et al., 2014; Thalhammer et al., 2024; Woodward, Karbasforoushan, & Heckers, 2012). Thalamic efferents were major contributors to the SZ-HC contrast, consistent with a disruption in thalamic gating and synchronization of cortical activity that may contribute to the intrusions and context- processing deficits characteristic of the disorder.

The posterior parietal cortex (PPC) is a core node of the frontoparietal working memory network, and its functional integration with lateral prefrontal cortex is crucial for working memory performance (Nielsen et al., 2017). In schizophrenia, PPC-prefrontal connectivity is consistently disrupted: dynamic causal modeling studies have shown reduced connectivity from inferior parietal lobe (IPL) to inferior frontal gyrus (IFG) during working memory tasks in first-episode patients (Nielsen et al., 2017), and parieto-prefrontal feedforward effective connectivity during context updating is reduced in patients with delusions, with the same connectivity adversely influenced by polygenic risk for schizophrenia in healthy individuals (Greenman et al., 2020). The present finding that PPC causal centrality is significantly lower in patients than controls is therefore coherent with this literature: rather than the PPC becoming pathologically dominant, its causal influence within the eDMN is attenuated. This reduction may reflect a failure of the PPC to exert its normal feedforward influence on prefrontal regions, consistent with the view that frontoparietal dysconnection—rather than parietal over-engagement—underlies working memory and cognitive deficits in schizophrenia.

Across these regions, a common thread emerges: those contributing most to the SZ-HC difference graph (VLPFC, PFC, BF, Tmp, thalamus) have documented roles in neuromodulation, executive control, and sensory/linguistic integration—hallmark domains of impairment in schizophrenia. The PPC fits this picture from the opposite direction: rather than contributing disproportionately to the difference graph, its causal centrality is significantly reduced in patients, consistent with attenuated parieto-prefrontal feedforward influence and frontoparietal dysconnection in the disorder. Together, these findings suggest that the altered causal architecture reflects a regionally specific redistribution of causal influence—temporal influence gaining prominence while parietal influence wanes— superimposed on a preserved core architecture in which subcortical regions remain causally dominant in both groups, rather than a uniform loss of function. The negative VLPFC and Tmp correlations with cognition further suggest that causal centrality in the eDMN is behaviorally meaningful, and that pathological reorganization, rather than compensatory recruitment, may underly the altered causal architecture—a hypothesis that future longitudinal or interventional work could test.

### Software note

A demo code for the computation of causal centrality is available at: https://github.com/tszarghami/CausalCentrality. SC-ICA is part of the Group ICA of fMRI Toolbox (GIFT): http://trendscenter.org/software/gift/. Spectral DCM, PEB and BMR have been implemented in SPM12: https://www.fil.ion.ucl.ac.uk/spm/. Graph-theoretical measures can be computed using the *centrality* function in MATLAB.

## Data availability

The data analyzed in this study were obtained from the COllaborative Informatics and Neuroimaging Suite Data Exchange tool (COINS; https://coins.trendscenter.org). Data collection was performed at the Mind Research Network, and funded by a Center of Biomedical Research Excellence (COBRE) grant 5P20RR021938/P20GM103472 from the NIH to Dr. Vince Calhoun. The improved parcellation map of the DMN (Alves et al., 2019) has been publicly shared by the authors on the NeuroVault repository: https://identifiers.org/neurovault.image:568084.

## Author contributions

TSZ: Conceptualization, Methodology, Analysis, Interpretation, Writing.

ORCID: Tahereh S. Zarghami: https://orcid.org/0000-0002-6208-2158

## Footnotes

1 Recent work has revealed the implication of DMN subsystems in the execution of certain external activities as well (see Mancuso et al. (2022) and the references therein).

2 https://identifiers.org/neurovault.image:568084

3 Generally, *θ* encompasses all the unobservable variables of the model, including the hidden states, parameters and hyperparameters. In this work, we only intervene on the causal influences (i.e., the effective connectivity parameters) of DCM, which shape the causal architecture of the graph.

4 Free energy is a lower bound on log model evidence, hence known as evidence lower bound (ELBO) in machine learning. In variational Bayesian inference, the optimized free energy serves as a proxy for log model evidence. Note that this variational free energy is the negative of free energy in statistical physics.

5 Here we make sure that the full model has been structurally optimized (using exploratory BMR); hence, intervention *decreases* the model evidence. Note that, for an over-parametrized model, edge (parameter) removal can *increase* the model evidence, which is the foundation of structure learning and optimization using Bayesian model reduction (Jafarian, Zeidman, Litvak, and Friston (2019); Beckers, van Erp, Zhao, Kondrashov, and Vries (2022); Neacsu, Mirza, Adams, and Friston (2022); Friston and Penny (2011)).

6 http://fcon_1000.projects.nitrc.org/indi/retro/cobre.html

7 Diagnostic and Statistical Manual of Mental Disorder, 4th edition (DSM-IV)

8 https://www.fil.ion.ucl.ac.uk/spm/software/spm12/

9 The Measurement and Treatment Research to Improve Cognition in Schizophrenia (MATRICS) initiative was designed to support the development of psychopharmacological agents to improve cognition in schizophrenia.

10 http://trendscenter.org/software/gift/

11 Note that vanilla spectral DCM is typically used for causal networks with less than 15 regions. For larger networks, further linearized variants of DCM can be used (Frässle et al. (2021); Friston et al. (2021))

12 Ventro-median prefrontal cortex (VMPFC)

13 Antero-median prefrontal cortex (AMPFC)

14 Dorsal prefrontal cortex (DPFC)

15 Temporal pole (TP)

16 Middle temporal gyrus (MTG)

17 Cerebellar hemisphere (CbH)

18 Cerebellar tonsil (CbT)

19 Since cross-spectra are the Fourier counterparts of cross-correlations, spectral DCM is essentially a causal model of how fMRI functional connectivity is generated.

20 Hence, the effective connections encoded in matrix *A* have units of *Hertz*. For more intuition, note that a differential equation of the form *x͘*(*t*) = *A* ∗ *x*(*t*), has a solution of the form *x*(*t*) ∝ exp(*A* ∗ *t*). As such, *A* admits units of 1/s = Hz.

21 Blood oxygenation level dependent (BOLD)

22 Self-connections control the region’s excitatory-inhibitory balance, or equivalently its gain or sensitivity to inputs.

23 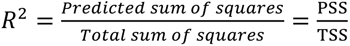, where PSS denotes the sum of squares of cross-spectra predicted/modeled by spectral DCM, and TSS denotes the total sum of squares of cross-spectra estimated from empirical data. The summations (of cross-spectra squared) are performed over frequency bins and pairs of regions for which the cross- spectra are computed. In SPM12, *R*^2^ for any fitted DCM can be calculated using spm_dcm_fmri_check.m.

24 Note that, the optimized model structure does not necessarily match the *ground truth*/*true model*. Because, by design, DCM offers the most accurate and least complex explanation for the given data, which may not match the more complex ground truth that has generated the data (Litvak et al. (2019)). As such, the modeling results and their interpretations are bound to the model assumptions and the model space.

